# A high-fidelity in vitro model of the human intestinal epithelium

**DOI:** 10.1101/2024.01.01.573838

**Authors:** Satoru Fujii, Scott T. Espenschied, Vibha Anand, Joao Bettencourt-Silva, Yi Han, Go Ito, Akira Koseki, Akihiro Kosugi, James Kozloski, Ryoma Matsumoto, Shanshan Meng, Natasha Mulligan, Ryan J. Musich, Kevin P. Newhall, Eri Ohshina, Shuhei Sekiguchi, Yi Wang, Jianying Hu, Matthew Ciorba, L. David Sibley, Ryuichi Okamoto, Thaddeus S. Stappenbeck

## Abstract

Primary intestinal epithelial stem cell culture methods have significantly advanced understanding of mammalian intestinal development and disease. However, progress has been hampered by inconsistent methodological reporting and challenges comparing *in vitro* systems with *in vivo* observations. We previously established a unique method for long-term 2­dimensional (2D) cultivation of mouse lECs using an air-liquid interface (ALI) technique which appears to faithfully recapitulate homeostatic and regenerative features in vitro. Here, we further refined these methods and optimized protocols for long-term, self-organizing 2D cultivation of human lECs. During the culture, we observed that epithelial cells undergo a dynamic morphological transition from squamous to columnar shape. Using single cell transcriptomics, we identified both major lineages and minor populations, including enteroendocrine cells, tuft cells, and BEST4/CA7+ cells. Leveraging the power and scalability of a biomedical foundation model (BMFM) trained on single cell RNA sequencing data, we performed classification tasks to identify cell types across sample sources and to quantitatively benchmark our in vitro differentiated cells against cells collected from patient biopsies. We observed a striking degree of similarity between our in vitro differentiated cells and the corresponding cell types in vivo for multiple differentiated lineages. This novel approach using BMFM holds promise to expand our understanding of the regulatory mechanisms including gene-gene regulation underlying homeostasis and regeneration as well as the functions of rare and poorly understood lineages within the human intestinal epithelia. Moreover, these methods are generalizable to other organs and can be used to assess the correspondence of cells across experimental modalities.

**Significance:** This manuscript addresses challenges in quantitative comparisons of cell types grown in vitro vs their in vivo counterparts. Here we use the intestinal epithelium as a model system to address this challenge. We devised an in vitro culture platform that supports multipotent intestinal epithelial stem cells and their numerous differentiated progeny. This system gives rise to all known rare and abundant lineages in correct proportions. Novel use of biomedical foundation models pre-trained on publicly available data and then fine-tuned to data from this platform enabled demonstration of high concordance of multiple in vitro differentiated lineages with the corresponding cells in vivo.

## INTRODUCTION

Continued advancement in understanding human intestinal development, physiology, and disease requires innovation in model systems. The ability to culture primary stem cells collected from intestinal biopsies was a significant vertical advancement in methods for experimental study of the intestinal epithelium (1). Guided by experimental results, subsequent technological iterations in intestinal epithelial stem cell (ISC) culture methods have identified conditions (including media and substrate/matrix compositions) which are permissive for differentiation of secretory and absorptive lineages. Air-liquid interface (ALI) cultures are a leading-edge technology for *in vitro* studies of mammalian epithelial tissues (2), but have not yet been optimized or applied to the human intestine. However, the diversity of experimental methods has created significant barriers to comparative– and meta-analyses. Coupled with often incomplete methodological reporting, this has effectively created a reproducibility crisis in the field of intestinal biology research.

The timeline of significant developments in ‘omics technologies has occurred roughly in parallel with major milestones in ISC culture methods, and have enabled robust genomic, epigenomic, and transcriptomic characterization studies of ISCs *in vitro* (3). Most recently, advances in single cell RNA sequencing (scRNAseq) have uncovered new insights into the diversity and putative functions of intestinal epithelial cells both *in vitro* and *in vivo* (4, 5). However, experimental validation confirming observations from scRNAseq studies has been scarce. Moreover, in spite of the plethora of scRNAseq data published over the last 5 years, computational methods for comparing multiple scRNAseq studies across different experimental modalities are limited, largely due to the lack of sufficient amount of high-quality labeled data, further hampering validation efforts.

Recently a new class of transformer-based machine learning algorithms, known as foundation models (6), has demonstrated remarkable capabilities to learn general representations (also called embeddings) from large amount of unlabeled data, which can then be used to create many task-specific machine learning models with dramatically reduced need for labeled data. While the most well-known demonstration of the power of foundation models have been through Large Language Models (LLMs) such as GPTx (7) and DeepSeek (8) recent studies has demonstrated the ability of such techniques to also ingest, process, and operate on large multimodal biological data classes, including single cell transcriptomics data (6, 9). By pre­training foundation models on scRNAseq data, and then fine tuning on project-specific tasks, there are potential analysis of single cell transcriptomics that were previously not feasible using traditional methods (6).

Here, we couple an advanced *in vitro* ISC culture system with single cell transcriptomics and Biomedical Foundation Models (BMFM) (http://research.ibm.com/proiects/biomedical-foundation-models). Using a variety of orthologous and complementary experimental methodologies, we demonstrate that our culture system faithfully recapitulates in vivo intestinal epithelial differentiation programs and lineage specification. We then leverage the power of BMFM to benchmark our *in vitro* intestinal epithelial model system against data from human biopsies by fine-tuning a pretrained BMFM single cell transcriptomics model (pre-trained on scRNAseq data set for the tasks of cell classification) (10). The resulting fine-tuned models were used to perform quantitative evaluation of similarities between *in vitro* differentiated intestinal epithelial lineages and putative concordant cell types present in patient biopsy samples using label transfer. We randomly split the experimental data for re-training or fine-tuning tasks into five sets for label transfer experiments.

## RESULTS

To define features of human intestinal epithelial cells (lECs) differentiated *in vitro, we* cultured human rectal epithelial stem cells from non-diseased donors *(SI Appendix* **Table 1)** first as spheroids enriched for epithelial stem and progenitor cells. From these cultures, we then established monolayers using a modified ALI culture technique **(Fig. 1A)** (11). An important difference from an orthologous mouse culture platform (11) was the requirement for including both TGFβ and ROCK small molecule antagonists during the establishment of both 3D and 2D (submerged) stem cell cultures, but both inhibitors are removed to permit differentiation under ALI conditions *(SI Appendix* **Fig. S1A).** Of note, while TGFβ blockade is not required for cultivation of mouse ISCs under maintenance or differentiation conditions, ROCK inhibitor was still required for culture maintenance of human and mouse ISCs *(SI Appendix* **Table S5).** While the addition of recombinant growth factors (EGF, FGF2, and IGF-1) was sufficient to induce proliferation of stem cells, these factors were not required for differentiation in the ALI platform *(SI Appendix* **Fig. S1A,B).** At ALI day 0 (ALIdO), the monolayer contained squamous cells with no obvious morphologic features of differentiation. By ALId7, the cells had uniformly adopted a columnar morphology, coincident with the emergence of goblet cell (GC)-like and enterocyte (EC)-like cellular features and resembling the homeostatic architecture of human rectum **(Fig. 1B).** Cell height, a surrogate for differentiation approached 40 µm **(Fig. 1C),** which is comparable to the IEC height observed in human colon tissues (12). (13). EdU incorporation assays revealed that the human IEC monolayer was hypo-proliferative at ALIdO, became transiently hyper-proliferative from ALId2-d4, then gradually returned to a less proliferative state thereafter **(Fig. 1D,E).** The transient hyper-proliferation mirrored our previous observations of the ALI culture of mouse lECs as well as regenerating rectal mucosa from ulcerative colitis patients, suggesting that the hyper-proliferative cells around ALId2-d4 may be reminiscent of regenerative processes in the human intestinal mucosa. Collectively, these observations focused our following studies on analysis of the ALI cultures at days 2 and 7.

**Fig. 1.**
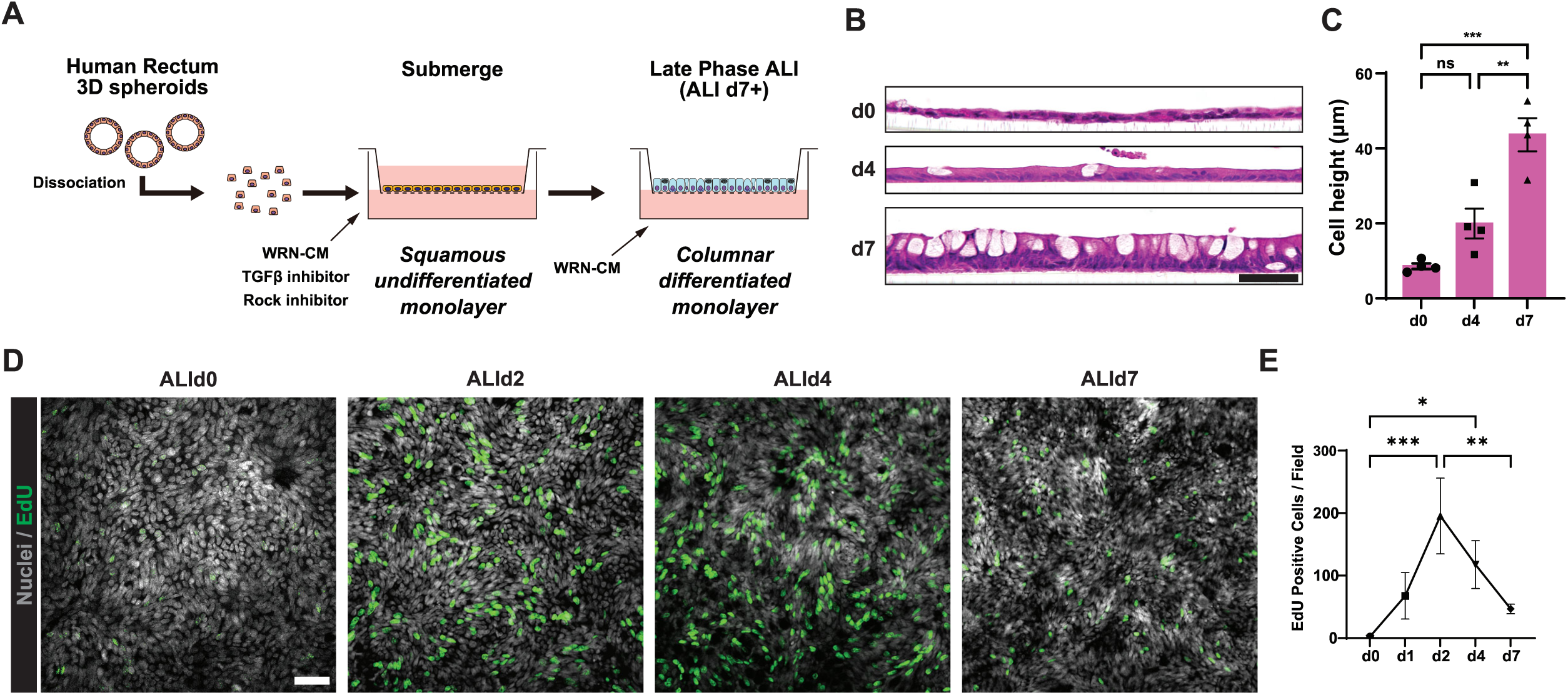
AU culture platform sustains differentiation of mature absorptive and secretory lECs. *(A)* Experimental schematic of ALI culture of human intestinal epithelial stem cells. *(B)* Hematoxylin and Eosin (H&E) images of human colon monolayer cells at days 0, 4, and 7 of ALI culture. Micrographs are representative of rectum lines established from 6 independent donors (n = 6); scale bar: 50 µm. (C) Epithelial cell height in ALI cultures at days 0, 4, and 7 (n = 4 rectum lines, data presented as mean ± SEM; significance determined ordinary one-way ANOVA with Tukey’s multiple comparisons test: **p < 0.005; ***p < 0.0005). *(D)* Representative fluorescence micrographs of EdU incorporation assay from ALI cultures at the indicated time points (nuclei in gray, EdU in green; scale bar = 50 µm). *(E)* Quantification of EdU^+^ cells (cultures established from n = 3 independent donors). In C and E significance was determined by ordinary one-wayANOVA with Tukey’s multiple comparisons test; *\*P* < 0.05, *\*\*P* < 0.005, *\*\*\*P* < 0.0005.

### Cross-platform comparison of scRNAseq from in vivo and ALI-cultured intestinal epithelial cells using traditional and novel methods

Given the temporal dynamics of human intestinal epithelial stem cell-derived cultures maintained under ALI conditions, we first used single-cell RNA sequencing to establish the transcriptomic heterogeneity of ALI-cultured human lECs using at ALId2 and ALId7 –. Following QC filtering *(SI Appendix* **Fig S2, Fig S3),** Uniform Manifold Approximation and Projection (UMAP) analysis revealed the presence of similar clusters in cultures established from 3 independent donors *(SI Appendix* **Fig. S3)** (14). Dimensionality reduction analysis of the integrated datasets revealed increasing separation of cell populations as the cultures matured **(Fig. 2A,B).** At ALId2 the cultures appeared transcriptionally homogenous, with generally poor separation of clusters in UMAP space. Manual cell typing using default analysis parameters was unable to confidently identify lineages – in spite of apparent clustering – at ALId2. However, by ALId7 manual cell-typing of the 11 clusters using canonical marker genes revealed the presence of major known cell types including: stem cells (SCs, Cluster #0), transit-amplifying cells (TAs, Cluster #1), enterocytes (ECs, Clusters #2, #3), goblet cells (GCs, Clusters #4-6), enteroendocrine cells (EECs, Cluster #9), and tuft cells (Cluster #10) **(Fig. 2B, Dataset_SOI).**

**Fig. 2.**
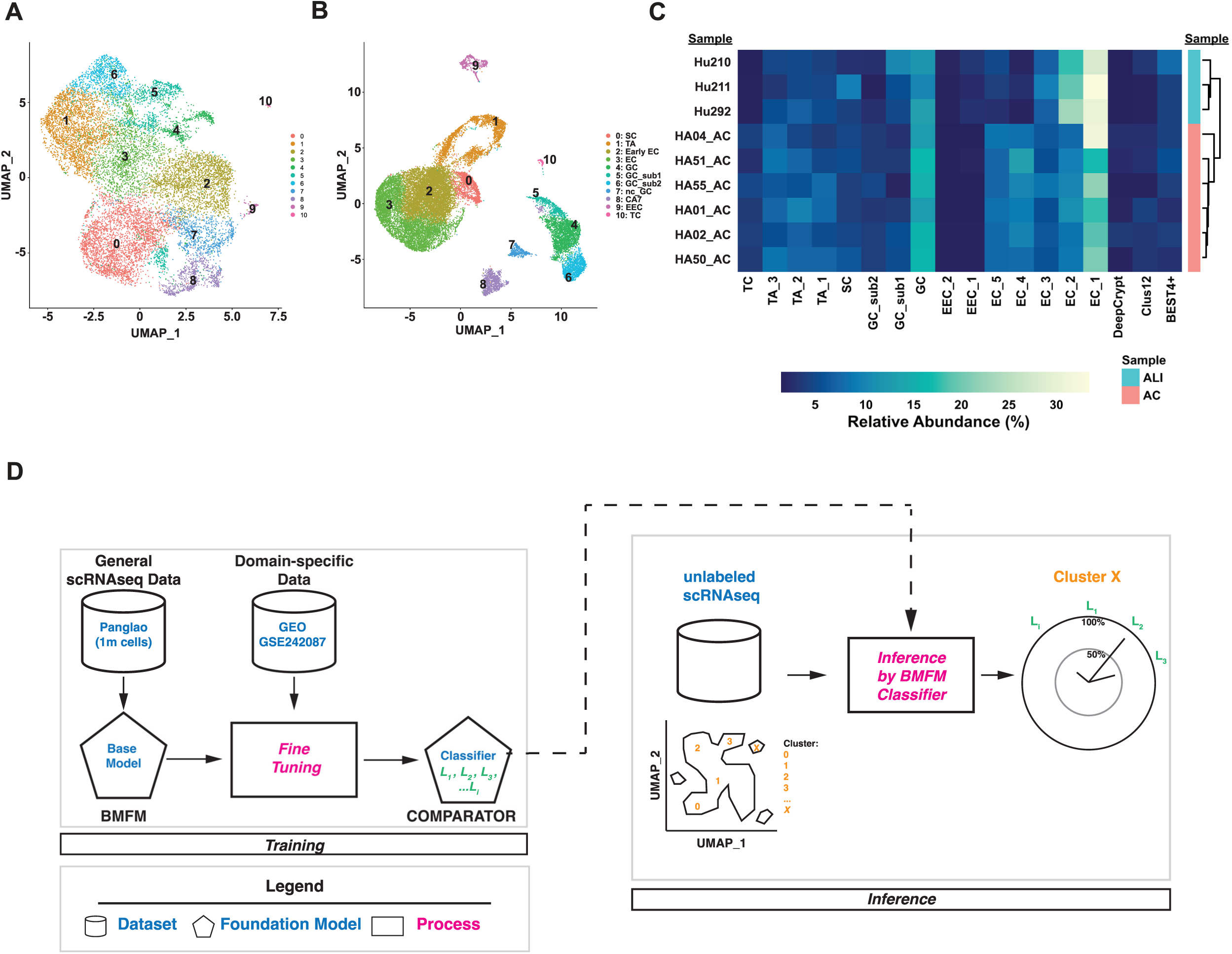
scRNAseq analysis of in vitro differentiated ISCs and classification of cell types using a domain-adapted Biomedical Foundation Model. (A,B) UMAP representations of integrated scRNAseq data from ALId2 (A) and ALId7(B). *(C)* Relative abundances of the indicated cell types from scRNAseq of ALI day 7 (n = 3) and human ascending colon (n = 6) *(D)* Schematic Biomedical Foundation Model Classifier (“COMPARATOR”) and *in silico* label transfer experiment.

Relative lineage abundances from the ALId7 scRNAseq data were similar when compared to the epithelial cell fraction from scRNAseq of ascending colon (AC) biopsies from 6 non-IBD patients *(X^2^* = 133.09, df = 136, *p* = 0.5546) **(Fig. 2C)** (10). These observations suggest the ALI culture platform is permissive for the differentiation and allocation of canonical absorptive and secretory lineages present in the human intestine. Based on these findings, we next utilized single cell data and Al methods to evaluate the degree of similarity of specific differentiated cell types in the colon. We hypothesized that we would find a high degree of statistical concordance between cells evaluated directly from patient biopsies with ALId7 cultured cells, further justifying the utility of this platform for modeling human cells and tissues.

Cross platform comparison and validation of cell types from different sources and platforms has been challenging since the inception of ‘omics approaches. To overcome this challenge, we designed a computational approach leveraging the power of foundation models: COMPARATOR (Computational Omics-based Modeling for Predictive Analysis of Response Alignment and Transfer Optimization in Real-world systems) enables cross-context translation between experimental systems and patient data through advanced label transfer, domain adaptation, and alignment verification technologies. Here ‘label transfer’ refers to the process of transferring labels from a pre-annotated, reference dataset to a new, unlabeled dataset that is the query **(Fig. 2D).** The goal is to use existing knowledge to efficiently annotate new data sets. Using one of the open-sourced BMFM models (https://qithub.com/BiomedSciAI/biomed-multi-omic) that was pre-trained for experimental modality-specific cell type annotation. We performed label transfer for all cells in all clusters in both datasets **(Fig. 2D, Dataset_SO2)** (10). We used radar plots as a visual representation of label transfer results (multivariate data).

Evaluation of BMFM classification results from ALId2 and ALId7 uncovered mixed labeling of many clusters at ALId2 (S/ *Appendix* **Table S6 and Table S7).** Consistent with the EdU incorporation assay, the clusters in the ALId2 samples appeared to be mostly comprised of proliferative TA-like cells. Interestingly, the BMFM label transfer experiment uncovered several groups of what appear to be ‘proto’ lineages (Clusters 3,4,5, and 9), which included mixed labels of ‘TA’ and a mature cell label (e.g. EC, GC), which may partly explain the inability to manually type many of the ALId2 clusters. As the classifier exhibited the strongest performance for identifying major and minor secretory lineages, we elected to focus on these cell types for further analysis.

### Goblet cells derived from the ALI platform are functionally and transcriptionally comparable to primary human colonic goblet cells

We found fidelity of GCs at ALId7 in the *in vitro* culture system with GCs in vivo both morphologically and by analysis of single cell transcriptomic data. Transmission electron microscopy (TEM) analysis of ALId7 cells showed ultrastructural features of GCs including a distinct and well-formed apical theca containing mucus granules **(Fig. 3A).** Targeted gene expression analysis showed enriched expression of the mucin-encoding gene *MUC2* at ALId7 at this time point **(Fig. 3B).** Immunofluorescence staining of ALI culture cross sections with an anti-MUC2 antibody indicated the presence of goblet cells with an overlying mucus layer **(Fig. 3C),** which was sufficient to prevent *Shigella* flexneri from directly contacting the I EC monolayer **(Fig. 3D).** These observations support that GCs in the ALI system are functional, and recapitulate protective mechanisms observed in mucosa *in vivo. \Ne* next assessed similarities between *in vitro* differentiated GCs and GCs identified in scRNAseq of biopsies from non-diseased patient donors. When we analyzed GCs from the reciprocal label transfer experiments. Label transfer to the immature ALI culture identified a cluster of proto-goblet cells defined by mixed TA and GC labels **(Fig. 3E).** However, by ALId7, we found similarly high labeling efficiency in both directions (O.93±O.OI7 from ALI to Biopsy; O.9O4±O.OO57, from Biopsy to ALI) **(Fig. 3F,G).** Thus, GCs from both data sets, by multiple observational and quantitative metrics, were highly similar to each other.

**Fig. 3.**
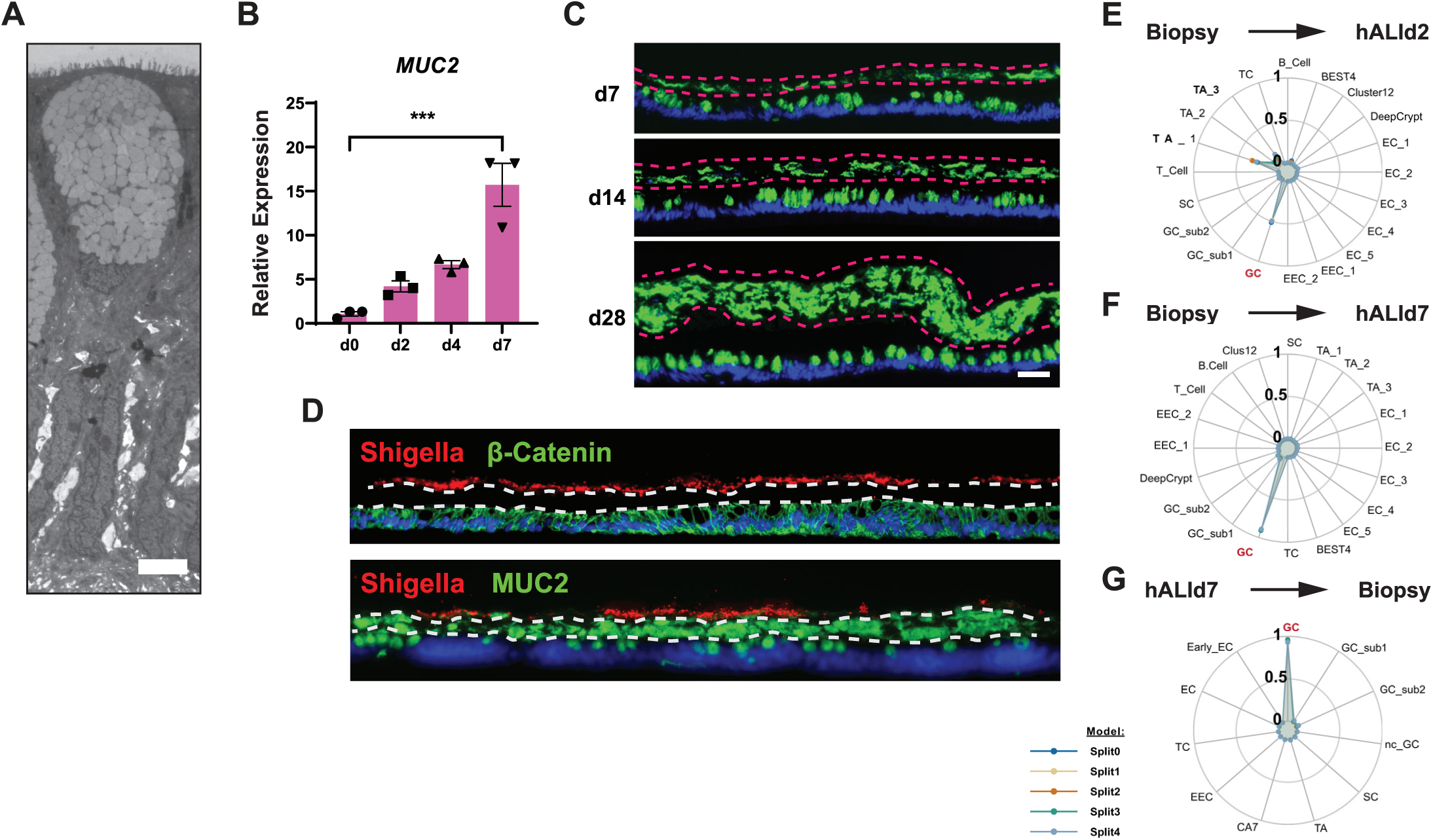
ALI culture supports differentiation of functional GCs. (A) TEM images showed the morphology of goblet cells at ALId7. Image is representative of 3 independent rectal lines. Scale bar: 5 µm. (B) qPCR analysis of *MUC2* expression in human rectum monolayers at ALI day 0, 2, 4 and 7. Results were normalized to the day 0, and shown as mean ± SEM (n = 3 rectum lines per time point). Ordinary one-way ANOVA multiple comparisons: *p < 0.05; **p < 0.005; ***p < 0.0005; ****p < 0.0001. (C) Representative fluorescence photomicrographs of ALI culture sections stained with an α-MUC2 antibody. *(D)* Immunofluorescence of *Shigella flexneri* (red) with beta-catenin (upper panel, green), and with MUC2 (lower panel, green), in homeostatic rectum monolayer cells during ALI cultivation, co­cultured with *Shigella flexneri* applied to the apical side of the monolayer. Representative images of three independent experiments. *(E-G)* Radar plot of BMFM label transfer experiment results for GCs (biopsy → hALId2, biopsy → hALId7; and hALId7 → biopsy)

### ALI culture enables differentiation of all known human rectal EEC subsets

We next turned our attention to hormone-producing secretory cells. Plotting gene expression in UMAP embedding space demonstrated enrichment of the EEC marker genes *CHGA* and *NEUR0D1* in Cluster 10 **(Fig. 4A,B).** Time course target gene expression analysis revealed relative CHGA expression peaked in the mature ALI culture at d7 **(Fig. 4C).** Amongst human lECs, enteroendocrine cells (EECs) are known to consist of various sub-populations, each of which produces distinct types of hormones (15). By isolating the EEC cluster, and performing sub-clustering, UMAP analysis we identified the presence of 5 discrete sub-populations of EECs in the ALId7 cultures **(Fig. 4D).** Further investigation of the ALI EEC scRNAseq cluster identified L-cells, which express GCG (glucagon) and *PYY,* enterochromaffin cells (ECCs), which express *TPH1,* and EEC progenitors, which express *NEUROG3* **(Fig. 4E,F).** Of note, a subset of EECs in the ALId7 monolayer cells forms a small sub-cluster that distinctively expresses *somatostatin* (SST), indicating the presence of D-cells **(Fig. 4E,F),** which have been observed in human colonic tissue sections. Previous scRNAseq studies failed to identify EEC subsets from human colon tissues or organoids (15–17). Immunofluorescence labeling with antibodies that specifically recognize NEUROG3, serotonin (an EC marker), and GLP-1 (an L-cell marker) in ALId7 cultures corroborated the scRNAseq analysis **(Fig. 4F).** We further confirmed the expression of SST at the protein level in mature ALI culture **(Fig. 4F).** Consistently, this finding corroborates the notion that the ALI culture platform supports *in vitro* differentiation of rare IEC secretory lineages found in the human colon.

**Fig. 4.**
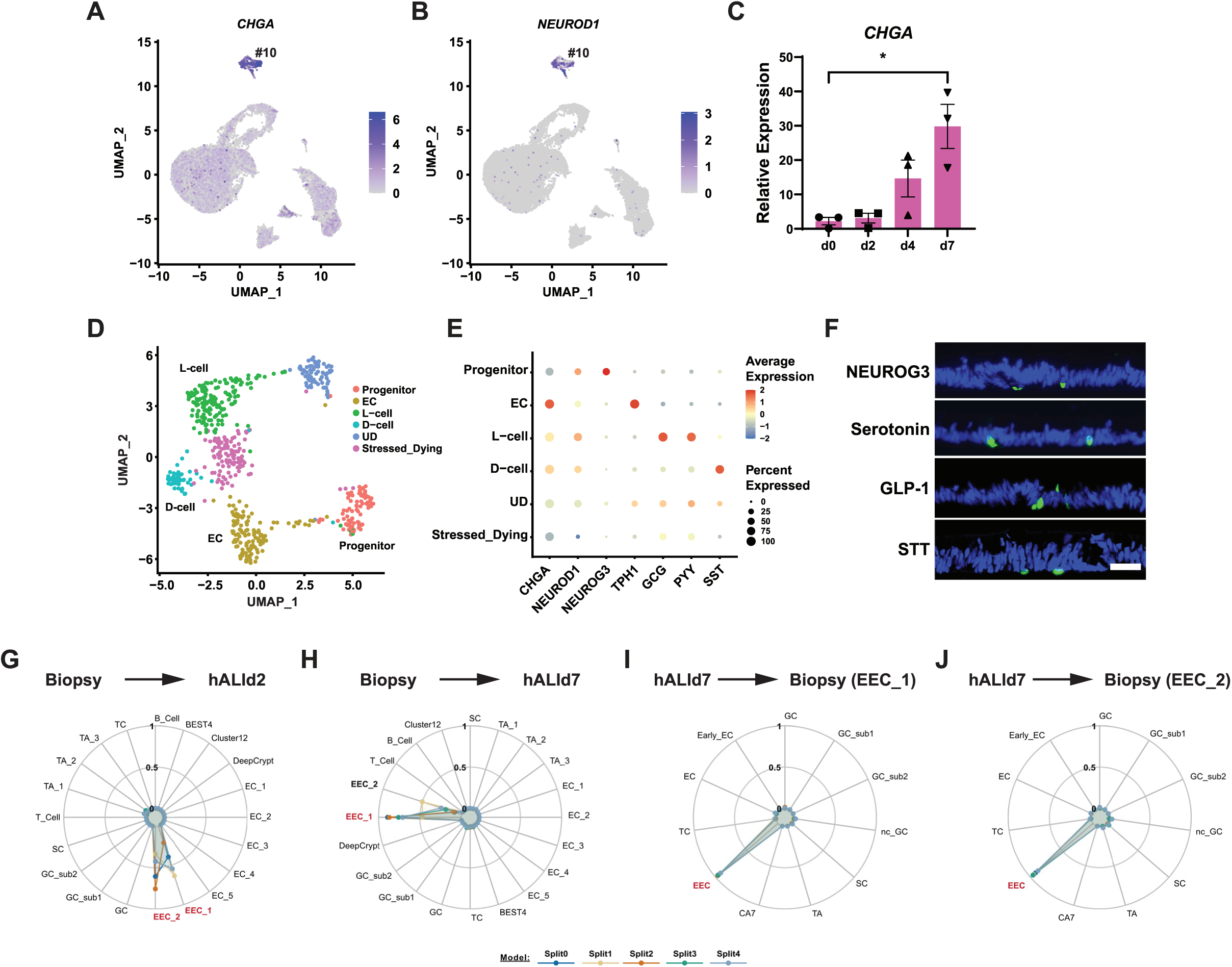
Differentiation of all known EEC-subtypes from rectal ISCs cultured under ALI conditions. (A,B) Feature plot of *CHGA* and *NEUROD1* expression in ALId7 cultures. (C) qPCR analysis of *CHGA* expression in human rectum monolayers at ALI days 0, 2, 4 and 7. Results were normalized to day 0. Values shown as mean ± SEM (n = 3 rectum lines per time point). Ordinary one-way ANOVA multiple comparisons: *p < 0.05; **p < 0.005; ***p < 0.0005; ****p < 0.0001. *(D)* Sub-clustering within the EEC cluster showing distinct sub-populations [Progenitor cells *(NEUROG3+),* ECs *(TPH1+),* L-cells (GCG+, *PYY+),* and D-cells *(SST+);* UD (undifferentiated, no specific EEC subcluster markers, Stressed_Dying cells have a higher mitochondrial gene ratio than the others]. (E) Dot plot analysis of sub-populations within the EEC cluster. *(F)* Immunofluorescence of NEUROG3, serotonin, GLP-1, and SST in the human rectum monolayer at ALI day 7. Representative images of three rectum lines (n =3). Scale bar: 50 µm. (G-/) Radar plots of label transfer experiment results for EECs (biopsy → hALId2, biopsy → hALId7; hALId7 → biopsy)

Leveraging the BMFM model for cell type classification, we again performed reciprocal label transfer experiments to assess the similarities between EECs in vitro and in vivo. Of note, we had identified 2 populations of EECs in the biopsy dataset without the need for additional sub­clustering. Using a BMFM fine-tuned on biopsy data, we observed nearly equivalent labeling of ALId2-Cluster10 as EEC_1 and EEC_2; however, at ALDId7 we found the majority of in vitro differentiated EECs were classified as EEC_1 (ũ.74±0.17), while almost one quarter were classified as EEC_2 (O.23±O.17); when the two labels were combined, the model classified 97% of *in vitro* EECs correctly **(Fig. 4G-I).** Label transfer in the opposite direction was comparably strong when evaluating biopsy EEC_1 cells (O.95±O.O2) or biopsy EEC_2 cells (O.94±O.O3) classified by a model fine-tuned on our *in vitro* data. Taken together, the results from transcriptional, morphological, and protein characterization of differentiated cells in the ALI system are highly concordant with primary colonic EECs.

### ALI in vitro Differentiated Tuft Cells are Transcriptionally Homologous to Tuft Cells in vivo

We next compared *in vitro* and *in vivo* differentiated tuft cells (TCs). Given the signals required for experimental expansion of TCs in mice (e.g., helminths, and IL25), we hypothesized that *in vitro* differentiated TCs in the ALI system would be rare and potentially transcriptionally distinct from TCs *in vivo,* as the former lacks requisite extrinsic signals required for the expansion and function (18, 19).

We identified a small cluster (Cluster #11) which exhibited distinctive expression of *TRPM5,* a putative marker for TCs in human lECs (20), assigning this to TC cluster **(Fig. 5A,B).** Notably, marker analysis identified 19 transcripts which are highly and uniquely expressed in TCs in the ALI culture **(Fig. 5A).** While *DCLK1* is the most common marker for detecting TCs in mice (21), no specific expression of *DCLK1* was observed in the TC cluster of the homeostatic monolayer cells **(Fig. 5C).** This is consistent with recent immunohistochemical and transcriptomic data suggesting that *DCLK1* is undetectable in human colonic tissues (22, 23), further supporting the notion that *DCLK1* does not identify TCs in humans. *PTGS1* (24), which encodes a critical enzyme in prostaglandin (PG) biosynthesis (25), and *POU2F3,* a transcription factor required for TC differentiation (26), both reliable alternative markers for human TCs, are distinctively expressed in this cluster (**Fig. 5B,** S/ *Appendix* **Fig. S5).** *SH2D6,* which has been recently identified as a marker for the rare sub-population of CD45^+^ TCs in mice, along with *PTPRC* (which encodes CD45) (27), are both enriched in this cluster **(Fig. 5B**, *SI Appendix* **Fig. S5A),** suggesting a potential subdivision of TCs in human intestinal mucosa, as observed in mice.

**Fig. 5.**
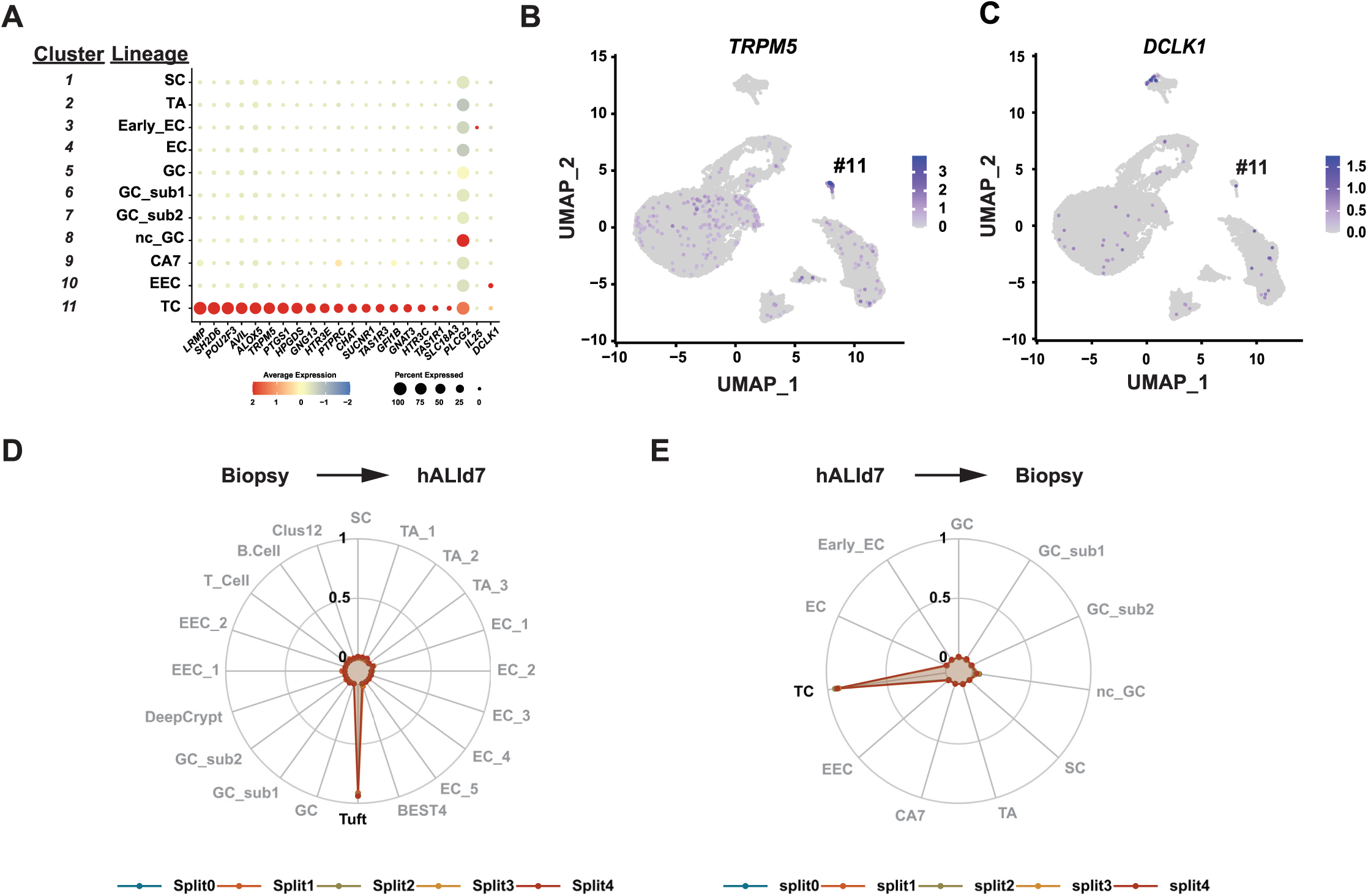
ALI culture permits spontaneous differentiation of chemosensory tuft cells. (A) Dot plot analysis of TC marker genes in ALId7 cultures. (B) UMAP feature plot of *TRPM5*. (C) UMAP feature plot of *DCLK1. (D-E)* Radar Plots of label transfer experiment results for TCs (left: biopsy → hALId7; right: hALId7 → biopsy)

Studies in mice suggest that TCs may be involved in the gut-brain axis and metabolic control, working in cooperation with enteroendocrine cells (EECs) and enteric neurons (21, 28). TCs express chemoreceptors, functioning as sentinels in the intestinal mucosa to regulate homeostasis and regeneration, as well as to respond to infections (18, 29). Intriguingly, pathway analysis on the TC cluster revealed that pathways related to serotonergic synapse and cholinergic synapse signaling are ranked as most prominent (S/ *Appendix* **Fig. S5B).** Further analyses identified specific expression of *CHAT,* coding for a critical enzyme for acetyl choline (ACh) synthesis (30), *SLC18A3,* which encodes a vesicular ACh transporter (31), and *HTR3C* and *HTR3E,* genes encoding serotonin receptors (32), in this cluster *(SI Appendix* **S5A).** The TC cluster also distinctively expressed the taste receptor genes *TAS1R1* and *TAS1R3 (SI Appendix* **Fig. S5A).** These findings corroborate the possibility that TCs in the human intestinal mucosa could function partly as neuronal transmitters/receptors, and/or chemoreceptors, as well as components of PG synthesis machinery. Of note, while studies in mice have clearly suggested that TCs function as a major source of IL-25 in the intestinal mucosa, crucial for initiating type 2 immune responses against helminths and protozoan parasites (18), the TC cluster cells did not exhibit specific expression of *IL25 (SI Appendix* **Fig. S45).** This implies that human TCs might possess little relevance to IL-25 production, or their expression of *IL25* could require activation through certain stimuli.

We also leveraged the BMFM label transfer approach to evaluate concordance between *in vitro* differentiated TCs and primary TCs sampled from human biopsy tissue. Of note, the foundation model failed to identify TCs in the ALId2 scRNAseq data. From reciprocal label transfer studies, we observed extremely high confidence in the model’s ability to classify TCs (Biopsy to ALI: O.932±O.OI; ALI to Biopsy: 0.919±0.01) **(Fig. 5D,E).** Thus, maintenance of extrinsic signals are not required for human ISCs to allocate TCs *in vitro* and for these TCs to differentiate with similar programs as *in vivo* TCs.

The functions of TCs in the human intestine remain largely unknown. However, some studies focusing on IBD have reported a reduction in TCs in the epithelial mucosa of IBD patients (22, 23), suggesting a potential association with IBD pathology. The future application of this novel 2D culture technique in researching human intestinal TC functions holds promise for enhancing our understanding of IBD pathophysiology.

### ALI culture model enables investigation of poorly understood CA7/BEST4 cells

Previously studies reported *BEST4^+^* cells as a specialized lineage of mature absorptive enterocytes in human intestinal tissues (33, 34). The *BEST4* gene, which encodes a Ca^2+^-sensitive chloride channel, has recently gained attention as *BEST4^+^* cells may be associated with pathophysiology of UC and colorectal cancer (17, 35). *BEST4^+^* cells co-express *OTOP2,* which encodes a proton-conducting ion channel, and both are thought to be engaged in fluid/mineral balance in human intestinal mucosa (17). In addition to *BEST4* and *OTOP2,* cells in Cluster #9 distinctly expressed the genes *CA7, HES4,* and *NOTCH2* **(Fig. 6A-C),** which were also reported to be expressed in the *BEST4^+^* cells *in vivo* (17, 36). We observed more uniform expression of *CA7* compared to *BEST4* in this cluster, prompting us to label this cluster “BEST4/CA7”. *In situ* hybridization analysis revealed a small but definitive number of CA7-positive cells in both ALI cells and human colonic tissues **(Fig. 6D),** confirming the findings from our scRNAseq analysis and demonstrating the existence of this minor population *in vitro.* We assessed similarities between the CA7^+^ cluster of cells differentiated *in vitro* under ALI conditions and BEST4^+^ cells in patient biopsies using the BMFM label transfer approach described above. Transferring labels to the ALId2 cultured cells, we observed heterogeneous labeling of ClusterX wherein ∼5O% of the cells were classified as BEST4, and a similar fraction identified as the enterocyte sub-type EC4 **(Fig. 6E).** At the later ALI time point, we found cells of the CA7 cluster were mostly labeled as BEST4^+^ cells [labeling efficiency of 0.93±ũ.017 (µ±σ)] when the model was fine-tuned on scRNAseq of epithelial cells from human biopsy samples. When we fine-tuned the model on our hALId7 scRNAseq data and applied labels to the non-IBD dataset, we observed less efficient labeling of the CA7 cluster as BEST+ cells (O.77±O.O25); which may be partially explained by the reduced cellular heterogeneity of our *in vitro* culture system **(Fig. 6F,G).**

**Fig 6.**
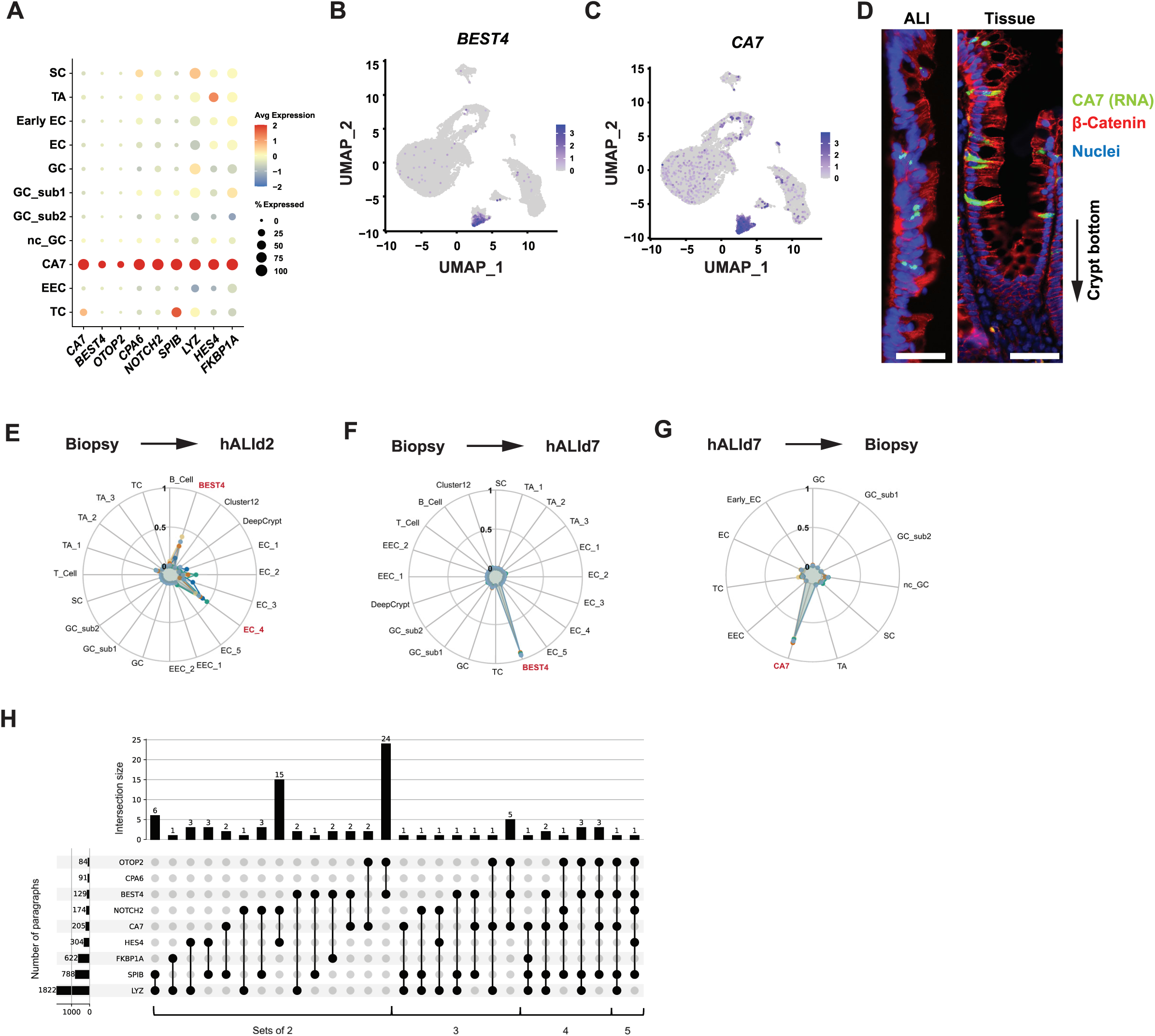
ALI culture platform enables investigation of CA7/BEST4+ cells. (A) Dotplot of marker genes for BEST4/CA7 cluster. *(B,C)* UMAP feature plots of *BEST4* and *CA7* expression. *(D)* Representative fluorescence photomicrographs of RNA *in situ* hybridization for *CA7* by RNAscope in the human rectum monolayer cells at ALI day 7 (upper panel) and human colon surgical resection tissue (lower panel) (n =3). Scale bar: 50 µm. *(E-G)* Radar plots of label transfer experiment results for BEST4/CA7 cells (left: biopsy → hALId7; right: hALId7 → biopsy). (H) UpSet plot of NLP literature search results for subsets of the BEST4/CA7 marker gene list.

The set of 9 marker genes which distinguish the BEST4/CA7 cluster is too short for functional pathway enrichment analysis but represents a combinatorially-intractable human literature search problem. We thus devised an *in-silico* approach using natural language processing to query the entire text of primary articles accessible from PubMed Central (see methods in S/ *Appendix* for details) for co-occurrence of subsets of marker genes for the BEST4/CA7 cells. This approach identified manuscripts in which pairs or higher-order subsets of genes (> 4 genes) from the CA7 marker gene list co-occurred in the same paragraph. Analysis of subsets of the marker genes revealed no co-occurrence of *CPA6* with any of the other genes. However, all higher-order subsets returned scRNAseq studies of the human gastrointestinal tract **(Fig. 6H, Dataset_O3)** (37, 38). Our search results returned no other organs or tissues, suggesting cells defined by this gene list are unique to the intestine.

## DISCUSSION

By integrating single cell RNA sequencing with the power of foundation models and classical molecular and histopathologic approaches, we dissected the features of *in vitro-* differentiated human intestinal epithelial cell lineages. scRNA seq of human rectal epithelial cells cultivated under ALI conditions identified 11 distinct clusters, including mature and functional absorptive and secretory cell lineages. While enterocytes and goblet cells are the most abundant cell types in mature cultures, we also found rare populations including TCs, EECs, and the poorly understood CA77BEST4^+^cells **(Fig. 1D).** Of note, the frequencies we observed in the ALI culture system are congruent with the frequencies of the same cell types detected by scRNAseq of biopsy samples from healthy donors. We were particularly encouraged by the presence of TCs in our culture system, as these cells are notoriously difficult to differentiate spontaneously *in vitro* (39, 40).

We leveraged the power of Biomedical Foundation Models to perform label transfer experiments by first training on scRNAseq from one dataset then performing cell type annotation of cells in a test dataset. Analysis of paired label transfer studies between biopsy and ALId7 datasets (i.e., “reciprocal label transfer experiments”) demonstrated the reliability of the foundation model classifier as well as the fidelity of the culture platform, as single cell transcriptional profiles of both abundant and rare epithelial cell lineages were highly concordant with cells recovered from biopsies **(Fig. 2E,F, 3G-I, 4D,E,** and **5D,E).** Similar label transfer analysis to the immature ALId2 cells revealed a variety of notable patterns, including co-labeling of clusters with “TA” and differentiated cell types, suggesting the presence of precursor or proto­cells. That said, not every cluster at ALId2 fit this pattern (e.g. EECs were clearly discernable on their own, while BEST4/CA7 cells imperfectly matched other cell types). Collectively, our findings demonstrate that the optimized ALI platform for culturing human intestinal epithelia faithfully recapitulates lineage allocation and differentiation of mature absorptive and secretory cell types which are highly concordant with cells observed *in vivo.* The methods reported herein describe an enabling technology which will facilitate the next generation of *in vitro* studies of the human intestinal epithelium.

Rigor and reproducibility are essential for scientific progress and maintaining public trust in the scientific enterprise. Here, we combine traditional experimental approaches with cutting­edge computational methods to thoroughly establish the fidelity of our new human intestinal epithelial stem cell culture platform. We corroborate expression of scRNAseq marker genes for GCs, EECs, and TCs by immunofluorescence labeling, *in situ* hybridization, and transmission electron microscopy. We further demonstrate GCs form a functional mucus barrier by co-culture with labeled bacteria. However, traditional validation methods are limited and do not scale well: only a few candidates can be tested at a time, and from a modest number of samples. Thus, there is a pressing need for highly dimensional and scalable unbiased data-driven techniques for quantitative comparison of scRNAseq. For example, using cell typing as a BMFM output will greatly expand the capacity for validation of scRNAseq studies. Moreover, these approaches may now be applied to experimental or in silico manipulations where lineage allocation is altered. These *in silico* experiments may subsequently inform the order, scope, or importance for predicting in vitro experimental outcomes.

Lastly, we applied a state-of-the art literature search tool (https://research.ibm.com/projects/deep-search) and natural language processing (NLP)-based tools to analyze the marker gene set which defines the CA7+/BEST4+ cells. This was done through iterative and combinatorial searching of the PubMed Central collection (up to April 2023). While we identified sets of 5/9 marker genes present in studies reporting scRNAseq analysis of human gastrointestinal samples, no manuscripts were identified with these gene sets in other organs or non-intestinal samples. However, our search space is currently limited to the main text of documents and does not include tabular data often appended to manuscripts as supplemental material. Thus, it is possible that cells with subsets of the CA7/BEST4 marker gene signature may exist in other organs. Experimental investigation of the ontogeny and function of these cells is an active area of investigation and will require the development of sophisticated cell biological systems in order to fully understand their properties. Ultimately, if applied during the planning stages of a new project, these tools may be further automated and combined with emerging Large Language Models (LLM) to assist researchers in streamlining future study plans by identifying gaps and interesting questions.

The challenge remains with scRNAseq of in vitro models to validate against in vivo results. Here we demonstrate a novel Al-enabled technology that when paired with a robust in vitro platform under differentiation conditions, identifies high degrees of concordance between cultured cells and their *in vivo* counterparts. Culture conditions without differentiation represent an opportunity to perceive an expansion of cell types which may play a role under pathological conditions, such as wound repair. Further use of these tools with other in vitro systems will enable cross-platform validation, thus increasing rigor and reproducibility of data coming from these systems.

## METHODS

### Human intestinal stem cell lines

Human stem cell cultures were established from rectal and ileal biopsies collected from non-IBD donors during routine endoscopy at the Washington University Digestive Diseases Research Center Core (DDRCC) or at the Tokyo Medical and Dental University Hospital. This study was approved by the Washington University Institutional Review Board, Cleveland Clinic Institutional Review Board), and the Tokyo Medical and Dental University Scientific Ethics Committee. Written informed consent was obtained from all donors. Isolated crypts were cultured as previously described (41). Demographic details of human donors are described in *SI Appendix* **Table S1.**

### Human Tissue

Formalin-fixed paraffin blocks of colonic surgical resection from non-IBD donors as healthy controls and UC patients were obtained from Department of Pathology and Immunology at the Washington University in St. Louis or Lerner Research Institute Biorepository Core at Cleveland Clinic. Sections (5 µm) were cut from blocks and stained with H&E, or processed for immunofluorescence or RNAscope *in situ* hybridization.

### Single cell RNA sequencing and analysis

Intestinal stem cell cultures were dissociated to single cell suspensions, purified by FACS (>70% viability, (>70% singlets), and immediately processed using the 10X Genomics Single Cell 3’ Reagents Kit v3.1 Next GEM protocol. Following quality control, libraries were sequenced on an Illumina Novaseq 6000 platform.

Sequence data were mapped to the human genome (GRCh38) using Cell Ranger (v7.0.1) with default para meters (42). Gene-barcode matrices were loaded in Seurat (v5.0.0) in R (v4.3.O)/RStudio (v2023.09.1+494) (43–47). Cells with high mitochondrial ratios (>20%) and/or high total transcript counts (>20,000) were excluded from further analysis. The top 3,000 most variable features were enumerated, and inter-subject variance removed via integration. Additional details are provided in the *SI Appendix*.

### Biomedical Foundation Models

IBM’s Biomedical Foundation Model for gene expression (biomed-multi-omic/biomed-gene-expression) was developed using the scBERT architecture (9) pre-trained on scRNAseq data from ∼1 million cells in the Panglao database (48) using transformer-based deep neural networks. After fine-tuning the model with scRNAseq data of either ALI cultured lECs or primary biopsy samples, the model was asked to classify known (to validate the model for cell type annotation) or unknown cells (to apply label transfer and establish cell type concordance).

## DATA AVAILABILITY

Single-cell RNAseq will be publicly available as of the date of publication.

## ACKNOWLEDGEMENTS

This work was supported by the Crohn’s & Colitis Foundation. We thank the Biorepository Core, Genomics Core, Flow Cytometry Core, and Imaging Core of the Cleveland Clinic Lerner Research Institute, and the Genome Technology Access Center and Digestive Disease Research Core Center at Washington University in Saint Louis.

## AUTHOR CONTRIBUTIONS

Conceptualization: TSS, SF, YW; Methodology: SF, AK, AK, JK, VA; Software: AK, AK, VA, JS, NM; Formal Analysis: STE, RJM, AK, JS; Investigation: SF, SM, Gl, RM, EO, SS, KPN, YH; Resources: MC, LDS; Data Curation: RJM; Writing – Original Draft: SF, STE, RO, TSS; Writing – Review & Editing: SF, STE, VA, JK, JS, JH, TSS; Visualization: SF, STE; Supervision: JH, TSS; Funding Acquisition: TSS

## Materials and Methods

### Crypt isolation and establishment of stem cell cultures

Human intestinal biopsies were minced with fine scissors and digested in 1 mL of 2 mg/ml_ collagenase type I solution (Gibco, 17100017) with 50 µg/ml_ gentamicin (Sigma-Aldrich, G1272) in washing media [DMEM/F12 (Sigma-Aldrich, D6421) supplemented with 1X L-glutamine (Gibco, G7513), 1X penicillin/streptomycin (CCF Lerner Research Institute Cell Culture and Media Core) and 10% FBS (Sigma-Aldrich, F2442)] for 20-30 min with pipetting every 5-10 min. Crypts were filtered through a 70-µm strainer, incubated with 9 mL of ice-cold washing media and pelleted by centrifugation at 300 x *g* for 5 min. Pelleted crypts were then suspended in 1 mL washing medium, transferred to a 1.5 mLtube and pelleted by centrifugation at 200 x *g* for 5 min. The supernatant was carefully removed by pipette, and the pellet was re­suspended in Matrigel (Corning, 354234). The Matrigel mixture was plated into 24-well tissue culture plates on ice. Plates were inverted and incubated at 37°C to polymerize the Matrigel and 500 µL of 50% L-WRN conditioned media (1) [a 1:1 mixture of L-WRN CM and fresh primary culture media (Advanced DMEM/F-12 (Gibco, 12634010) supplemented with 20% FBS, 1x GlutaMax (Gibco, 35050061), 1x penicillin/streptomycin)], supplemented with 10 µM ROCK inhibitor (Y-27632) and 10 µM TGF-β inhibitor (SB 431542 or A83-01). Media was refreshed every 2-3 days. Rectal stem cells grown as spheroid cultures were collected in Cell Recovery Solution (Corning, 354253), digested with TrypLE Express (Gibco, 12605010) into small fragments, and passaging every 3-4 days. These human rectal spheroid lines were maintained and passed up to 30 generations for the experiments reported in this paper.

### Air Liquid Interface (ALI) culture

Grown human rectal spheroids were dissociated into single cells by TrypLE Express digestion and seeded onto transwell membranes (0.4 µm pore size, Corning, 3470) pre-coated with 10% Matrigel diluted in PBS for 2Omin. 200,000 cells were seeded per well. Initially, 150 µL and 350 µL of 50% L-WRN media with 10 µM ROCK inhibitor (Y-27632) and TGF-β inhibitor (SB 431542 or A83-01) were added into the upper and lower chambers, respectively, and the cultures were maintained under submerged conditions for 5-7 days until achieving confluency. Medium in the upper chamber was then removed to create an air-liquid interface. Cells were grown in ALI with 50% L-WRN medium (with neither ROCK inhibitor nor TGF-β inhibitor) in the bottom chamber. Cells were cultured for an additional 7-28 days to reach maturity, and media changed every 2-3 days. Details regarding small molecules and recombinant peptides can be found in **Supplementary Table S2.**

### Histology

ALI cultured cells were fixed in 4% paraformaldehyde (PFA) for 30 min at room temperature. The fixed cells were then dehydrated in 70% ethanol and were embedded in 2% agar followed by paraffin embedding, sectioning, and staining. Bright field histological images were acquired with an Olympus BX61 microscope equipped with a DP22 camera and cellSens Standard v3.2 software. Cell height was quantified by averaging measurements from two slides per time point for each donor sample using cellSens Imaging software.

### Immunostaining

Slides were de-paraffinized and subjected to heat-induced antigen retrieval in Trilogy solution for 2Omin. Slides were blocked in PBS containing 2% BSA, 5% serum and 0.1% Triton-X for 1 hr at room temperature, and then treated with primary antibodies at desired dilutions overnight at 4°C. They were then washed 3 times in PBS and incubated with fluorophore-conjugated secondary antibodies for 1 h at room temperature. Fluorescent images were acquired with a Zeiss Axiovert 200M or a Zeiss AxioObserver 7 inverted microscope equipped with a Colibri 7 Light Source and Axiocam 702 detector. FIJI (Imaged) (v1,54f) was used to uniformly adjust brightness and contrast. Information about primary antibodies and dilutions can be found in **Supplementary Table S3.**

### Transmission electron microscopy

Cultures grown on transwell inserts were washed once in PBS, then fixed overnight at 4°C in 2.5% Glutaraldehyde (Electron Microscopy Sciences, 16120) and 4% Paraformaldehyde (Electron Microscopy Sciences, 15710) in O.2M Sodium Cacodylate Buffer (pH 7.4, Electron Microscopy Sciences, 11652). Fixative was removed and samples were stored in O.2M Sodium Cacodylate Buffer at 4°C until further processing. Transwell membranes were removed from inserts, cut in half, then embedded in Eponate 12 Resin (Pelco, 18012). Sections were cut with a Leica EM UC7 ultramicrotome. Semi-thin sections were transferred to glass microscope slides, stained with toluidine blue and imaged with an Olympus BX43 microscope equipped with a DP22 camera using 2Ox (NA 0.50, WD 2.10 mm), 4Ox (NA 0.75, WD 0.51 mm), and 6Ox (NA 1.25, WD 0.12 mm) objectives. Thin (200 nm) sections were stained with osmium tetraoxide and uranyl acetate, transferred to copper grids, and imaged using an FEI Tecnai G2 Spirit BioTWIN transmission electron microscope equipped with an 11-megapixel Orius 832 CCD camera. Operating voltages were held constant within imaging sessions.

### ATP assay

Human rectum 3D spheroids cultured in 50% L-WRN conditioned media with ROCK inhibitor and TGF-beta inhibitor were dissociated into small fragments and mixed with Matrigel at a concentration of 1000 cells/µL. The cell mixture was seeded onto a white opaque 96 well plate at 5 µL per well (5000 cells per well), and 100 µL of test media was overlaid per well. After 72 hours of incubation, an ATP assay was performed using the CellTiter-Glo 3D Cell Viability Assay (Promega, G9682), according to the manufacturer’s protocol. Luminescence was measured using a Promega GloMax Discoverer Microplate Reader. Viability was calculated according to the following equations:

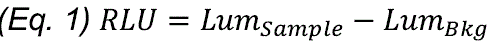

Where RLU is Relative (corrected) Luminescence, *Lum_samp_ι_e_* is the luminescence value measured in sample wells, and *Lum_Bkg_* is background luminescence values from wells with media only (no cells).

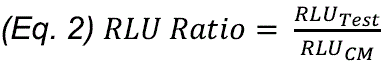

Where *RLUcm* is the Relative Luminescence of cells grown in 50% L-WRN conditioned media and *RLU_test_* is the Relative Luminescence of cells grown under experimental (“test”) conditions.

### Quantitative RT-PCR

RNA was extracted from ALI cells using the RNeasy Micro Kit according to the manufacturer’s protocol. Briefly, ALI cells were harvested in RLT buffer with 2-mercaptoethanol and stored in –80°C until use. RNA was extracted and eluted in 14 µl of nuclease free water and stored at –80°C until use. RNA was quantified on Cytation 5 by diluting 1:1 with nuclease free water to determine the volume of RNA needed to convert to cDNA. cDNA was prepared using 1 µg of starting RNA and reverse transcribed using iScript reverse transcriptase kit (Bio-Rad, 708841) according to manufacturer’s directions. Quantitative PCR reaction was conducted with TB Green Advantage qPCR 2X-premix (Takara, 639676) using a Bio-Rad CFX Connect Real-Time PCR Detection System. Primers were validated for specificity using a water control and conducting melt curve analysis. Data were analyzed using the Delta-Delta Ct method. Primer sequences for quantitative PCR are listed in **Supplemental Table S4.**

### RNAscope based in situ hybridization

ALI culture cells were fixed in 4% PFAfor 30 min at room temperature and sequentially cryoprotected in 10% (w/v), 15% (w/v), and 20% (w/v) sucrose in PBS, and frozen in Tissue-Tek OCT compound (Sakura, 4583). Cryosections (5 µm thick) were used for RNAscope-based *in situ* hybridization according to the manufacturer’s recommended protocols for RNAscope Multiplex Fluorescent Reagent Kit v2 (ACDBio, 323100). The probes Hs-LGR5 (ACDBio, 311021) and Hs-CA7 (ACDBio, 431961) were used in this study.

### Sample preparation for scRNAseq

ALI transwells were washed once with 1X DPBS-EDTA (O.O5mM) and incubated with TrypLE Express Enzyme at 37 °C for 25-30 mins, then pipetted 10 times with P1000 pipette and collected into a 15ml conical tube. Cell suspensions were incubated at 37°C until the cells mostly dissociate into singlets. The dissociated cell suspension was filtered through 20 µm strainer to purify cell singlets. A Denovix CellDrop FL automated cell counter was used to evaluate viability and singlet percentage of the prepared cell singlets. If necessary, the prepared cell singlets were sorted with BD FACS Melody Flow Cytometer by using Propidium Iodide and DRAQ5 double staining. The prepared cell singlets from ALI (viability>7O%, singlet percentage>7O%) were immediately processed according to 10X Single Cell 3’ Reagent Kits V3.1 Next GEM protocol (document # CG000204). Initially, 10,000 single cells were targeted using the Chromium Controller for cDNA synthesis and barcoding. Subsequent steps involved evaluating cDNA quality and quantity with a Bioanalyzer High Sensitivity DNA assay. The cDNA was then used as input for fragmentation, end-repair, adapter ligation, and sample indexing. Bioanalyzer assays ensured proper library construction. Libraries were pooled, quantified using a Quantabio Q cycler, denatured, and sequenced on an Illumina Novaseq 6000 platform. The sequencing protocol comprised 28 cycles for the forward read and 91 cycles for the reverse read.

### scRNAseq data processing

Sequence data were mapped onto the human genome (GRCh38) using CellRanger (v7.0.1) with default parameters. Fordownstream analysis, filtered gene-barcode matrices generated by CellRanger were read into the Seurat package (v5.0.0) on R (v4.3.O)/RStudio (v2023.09.1+494) (2). Plots were generated using RStudio and ggplot2, and further edited (for size and readability) with Adobe Illustrator CC. In Seurat, dead cells with high mitochondrial ratios (>20%) and potential doublets with high total transcript counts (>20,000) were first filtered out. The resulting data were normalized using SCTransform with default parameters. The top 3,000 most variable features were calculated, and integration was performed to remove patient-to-patient variance for clustering. The first 40 PCs were used for graph-based clustering, which was visualized with UMAP-based dimensional reduction (3). Marker genes for each cluster were calculated using the wilcoxauc function from the presto package (v1.0.0). Visualization of gene expression was performed using various functions within the Seurat package (i.e., FeaturePlot, DoHeatmap, and DotPlot).

### Biomedical Foundation Model: Label Transfer Experiments

Biomedical Foundation Models for Targets (BMFM) were developed at IBM research and pre-trained on public datasets using state-of-the-art deep neural network architectures based on BERT models (4, 5). In general, BMFM family of models capture broad knowledge across the biomedical domain and have been open-sourced by IBM (https://qithub.com/BiomedSciAI) (6, 7). BMFM uses attention mechanisms for learning large-scale, context-specific modeling (8). As transformers, these models are particularly well-suited to understand gene network dynamics as they provide universal representations of knowledge regarding gene expression derived from scRNAseq. Application specific fine-tuning using either the scRNAseq data generated in this study or the data from Zhao et al., enabled both validation of the cell-type annotation task as well as classification of unknown cells (9). Given an input dataset, the model can classify each cell by assigning labels from the fine-tuning dataset and generating a probability score (0–1) for each assignment; subsequently, the label with the greatest probability score is used as the predicted cell-type classification.

For the reciprocal label transfer experiment, the pre-trained BMFM was fine-tuned with either i) the hALId7 scRNAseq dataset (BMFM-hALId7) or ii) the epithelial fraction of both terminal ileum and ascending colon from the non-IBD controls in Zhao et al. (BMFM-ZhaoCtrlEpi) (9). Labels in the datasets used for fine-tuning were generated by manual curation of Seurat outputs informed from historical wet-lab experiments that defined marker genes for different intestinal epithelial cell lineages. This training methodology is schematized in *SI Appendix* **Fig. S5.**

Label transfer experiments were carried out 5 times. For each fine-tuned model, we randomly sampled 80% of the data from the fine-tuning set, 10% of the data for internal test and 10% of the data for internal validation. Label probabilities of each classification from all 5 fine­tuned models were plotted in R using ggplot2 and ggradar packages.

### Literature Gene Search and Analysis with Natural Language Processing

Document identification was performed using the Deep Search toolkit (10) and the full PubMed Central collection up to April 2023 was searched. All articles mentioning either one of the genes identified in the CA7/BEST4 cluster were selected and in the case of NOTCH2*, due to the very large volume of documents (N=5,I53), only combinations of this gene with any of the others were considered (N=160). Filters were applied to only keep genes that were identified by an automatic biomedical annotator (11) and that were mentioned in paragraphs in the main article text. The general workflow is illustrated in *SI Appendix* **Fig. S6.** Combinatorial results by gene subset were visualized using UpSet plots (12).

### ALI co-culture with Shigella flexneri!

*Shigella flexneri* Castellani and Chalmers (M9OT Sm) was cultured in TSB media to an optical density OD_6_qo = approximately 1.0. The bacterial culture was pelleted by centrifugation at 3500 x *g* for 3 minutes, after which the bacterial cells were resuspended in 2 mL of advanced DMEM with 20 mM HEPES. Bacterial suspension (100 µL) was applied to ALIdl 9 monolayer cells at a density of 500 bacterial per transwell. The co-culture was centrifuged for 5 min at 500 x *g,* after which the bacterial suspension was aspirated. After 24 hours of co-culture, the cells were fixed and processed for immunostaining and imaging.

### Statistical analysis, plotting, and data visualization

Statistical analyses and plotting were performed using Graphpad Prism (v9) and R/Rstudio (build/v). Statistical tests are reported in figure legends. A Mann-Whitney µ-test was used for two-sample comparisons, and ANOVA with Tukey’s multiple comparison’s test was used for comparing >2 groups. A *P* value < 0.05 was considered statistically significant. For comparison of lineage frequencies **(Fig. 1E),** a *X^2^* test was used.

## Tables

**Supplementary Table S1:** Human subject characteristics.

| Number | Sex | Age | Race | Location | Diagnosis |
| --- | --- | --- | --- | --- | --- |
| 1 | Male | 61 | Caucasian | Rectum | Healthy Donor |
| 2 | Female | 55 | African | Rectum | Healthy Donor |
| 3 | Female | 57 | Caucasian | Rectum, ileum | Healthy Donor |
| 4 | Female | 70 | Caucasian | Rectum | Healthy Donor |
| 5 | Female | 56 | Caucasian | Rectum, ileum | Healthy Donor |
| 6 | Female | 67 | Caucasian | Ileum | Healthy Donor |
| 7 |  |  | Asian |  | Healthy Donor |

**Supplementary Table S2:** Chemicals, Peptides, and other Reagents.

| Reagent | Vendor | Catalog_Number | Use |
| --- | --- | --- | --- |
| Y-27632 dihydrochloride | R&D Systems | 1254 | ROCK inhibitor |
| SB 431542 | R&D Systems | 1614 | TGFβ inhibitor |
| A83-01 | Tocris | 2939 | TGFβ inhibitor |
| EGF, mouse recombinant | Peprtech | 315-09 | growth factor |
| IGF-1, mouse recombinant | BioLegend | 590904 | growth factor |
| FGF2, human recombinant | Peprtech | 100-18B | growth factor |

**Supplementary Table S3:** Antibodies.

| Target | Host | Conjugate | Vendor | Catalog_Number | Clone | Application | Working_Concentration | RRID |
| --- | --- | --- | --- | --- | --- | --- | --- | --- |
| MUC2 | Mouse | N/A | Santa Cruz Biotechnologies | sc-515032 | F-2 | immunofluorescence | 1:100 | AB_2815005 |
| Chromogranin A | Rabbit | N/A | Abcam | ab15160 | polyclonal | immunofluorescence | 1:2000 | AB_301704 |
| 5-HT | Goat | N/A | ImmunoStar | 20079 | polyclonal | immunofluorescence | 1:5000 | AB_10719366 |
| GLP-1 | Rabbit | N/A | Bioss | BS-0038R | polyclonal | immunofluorescence | 1:200 | AB_10855332 |
| Neurogenin 3 | Rabbit | N/A | Bioss | BS-0922R | polyclonal | immunofluorescence | 1:200 | AB_10856983 |
| SST | Rabbit | N/A | Sigma Aldrich | HPA019472 | polyclonal | immunofluorescence | 1:1000 | AB_1857360 |
| β-Catenin | Mouse | N/A | BD Bioscience | 610153 | Clone 14/Beta-Catenin | immunofluorescence | 1:1000 | AB_397554 |
| Shigella | Rabbit | N/A | Abcam | ab65282 | polyclonal | immunofluorescence | 1:1000 | AB_1142846 |
| Rabbit | Donkey | AlexaFluor 488 | ThermoFisher | A-21206 | polyclonal | immunofluorescence | 1:200 | AB_2535792 |
| Mouse | Donkey | AlexaFluor 488 | ThermoFisher | A-21202 | polyclonal | immunofluorescence | 1:200 | AB_141607 |
| Goat | Donkey | AlexaFluor 488 | ThermoFisher | A-11055 | polyclonal | immunofluorescence | 1:200 | AB_2534102 |
| Rabbit | Donkey | AlexaFluor 594 | ThermoFisher | A-21207 | polyclonal | immunofluorescence | 1:200 | AB_141637 |

**Supplementary Table S4:**
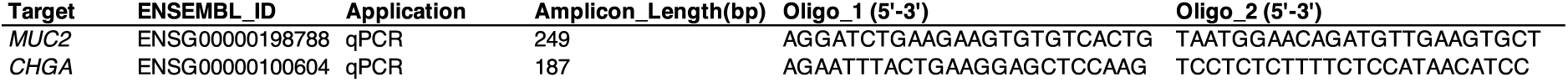
Oligonucleotides.

*Supplementary Table S5: Summary and Quality Control data for scRNAseq*

## Datasets

*Dataset_S0I: Marker gene sets for 11 clusters in differentiated ALI cultures*

*Dataset_S02: Label Transfer Experiment Results*

*Dataset_S03: DeepSearch Literature Search and Analysis Query of BEST4/CA7 marker gene signature*

## Resutls

**Fig. S1:**
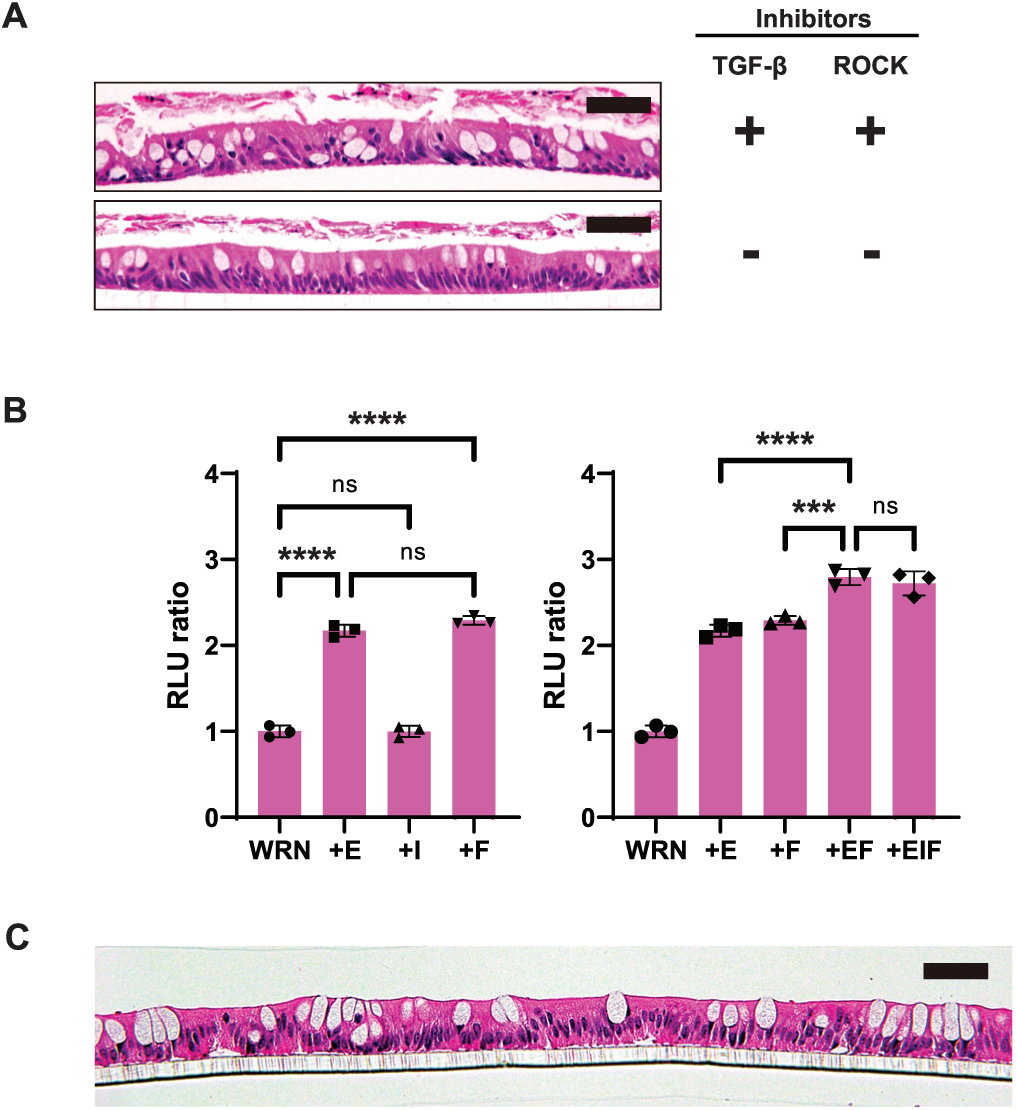
Optimization of culture conditions for differentiation of human ISCs in ALI platform. (A) Brightfield photomicrographs of H&E-stained sections of human rectal ISCs differentiated under ALI conditions in the presence/absence of TBF-β blockade (B) Effects of various growth factors on human ISC viability (C) Brightfield photomicrograph of H&E-stained section of human ileal intestinal stem cells differentiated under ALI conditions

**Supplementary Table S5:** Small molecule inhibitors required for mouse and human ISC culture under various formats.

| **Format** | **3D (Matrigel)** |  | **2D (submerged)** |  | **2D (ALI)** |  |
| --- | --- | --- | --- | --- | --- | --- |
| **Inhibitor** | **TGF-β** | **ROCK** | **TGF-β** | **ROCK** | **TGF-β** | **ROCK** |
| *Mouse* | - | - | - | + | - | - |
| *Human* | + | + | + | + | - | - |
| *Shared?* | No | No | No | Yes | Yes | Yes |

**Figure S2.**
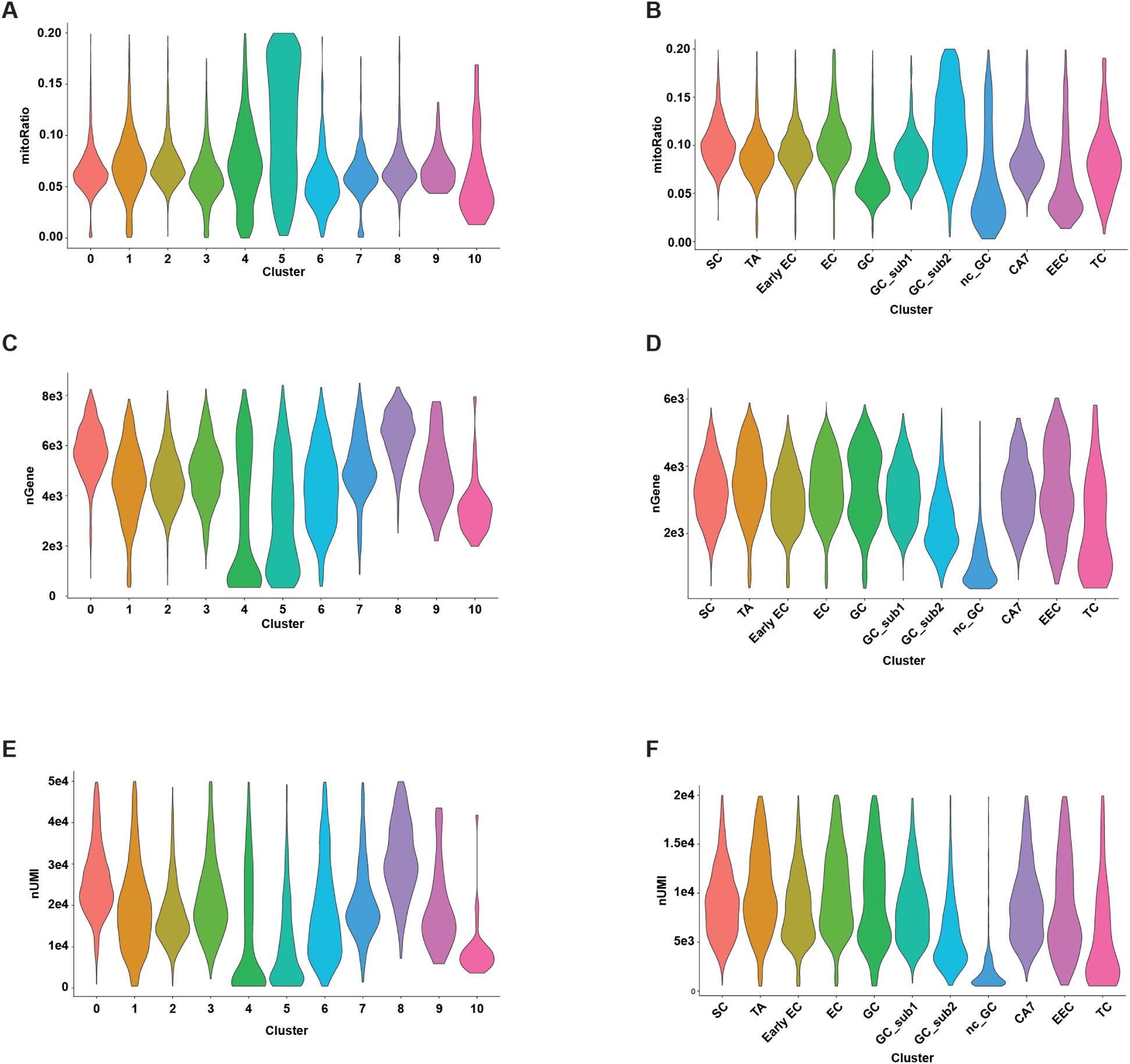
scRNAseq QC analysis by cluster. (A,B) Violin plots of mitochondrial ratio (mitoRatio) split by cluster from ALId2 and ALId7 samples. (C,D) Violin plots of number of genes (nGene) per cell split by cluster from ALId2 and ALId7 samples. (E,F) Violin plots of number of unique molecular identifiers (nUMI) per cell split by cluster from ALId2 and ALId7 samples.

**Figure S3:**
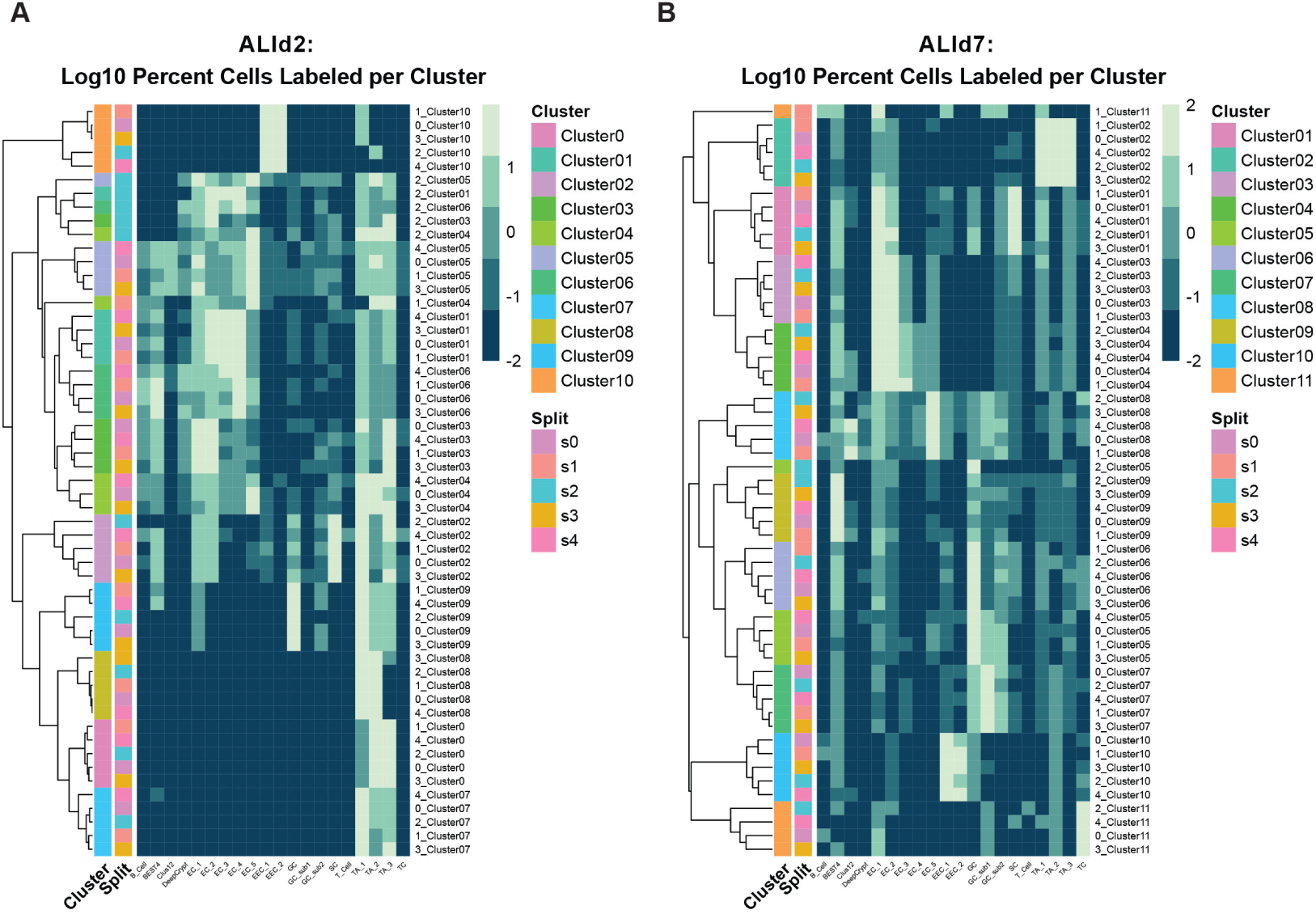
Summary of label transfer results: *in vivo* to *in vitro*. (A) ALI day 2 labelling frequencies by UMAP cluster (B)ALI day 7 labeling frequencies by UMAP cluster

**Fig. S4.**
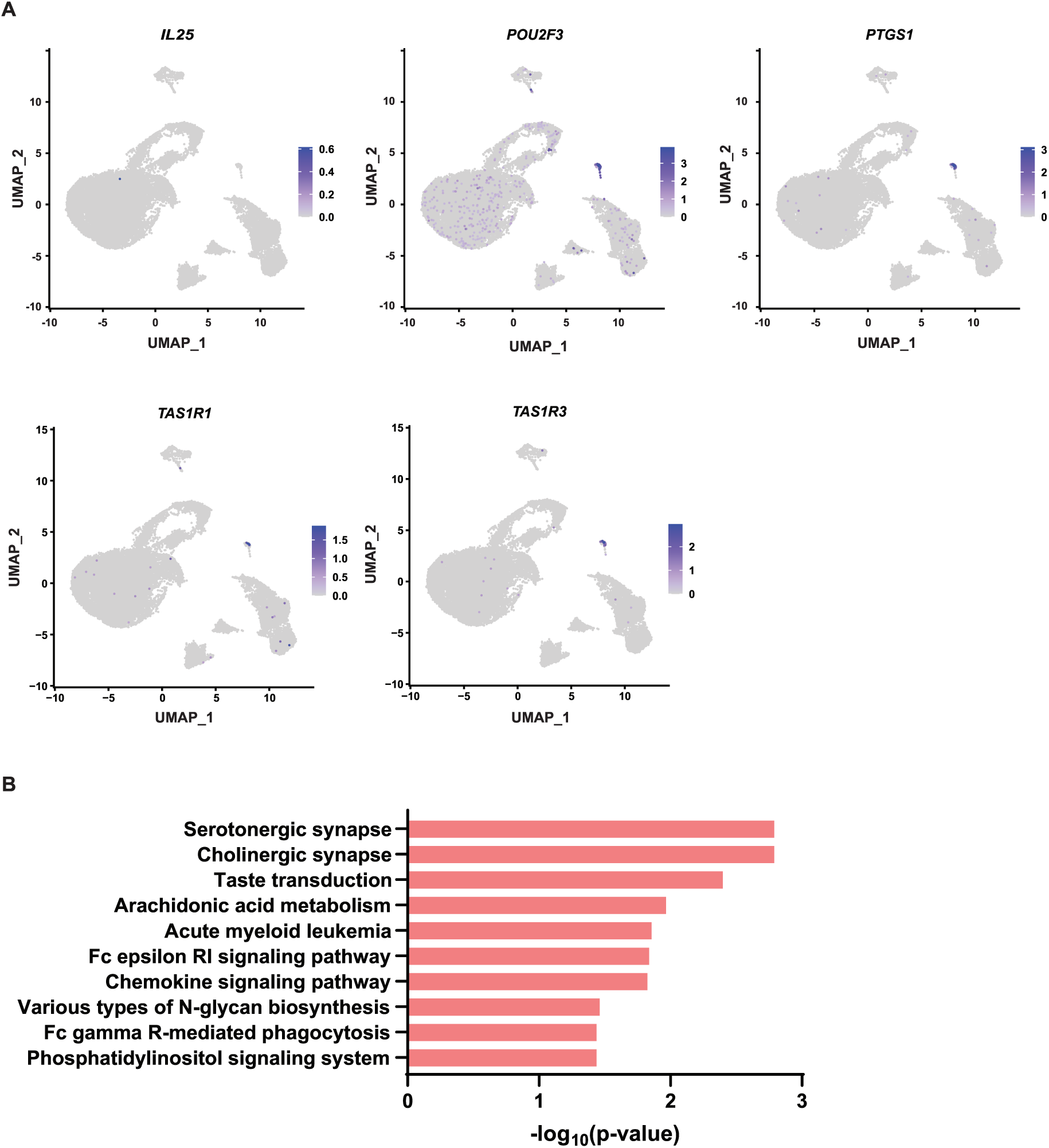
Expression of canonical and non-canonical TC marker genes in ALI-differentiated human ISC cultures. (A) Feature plots of TC marker gene expression in UMAP space (B) KEGG Pathway enrichment analysis of top 150 TC genes

**Fig S5.**
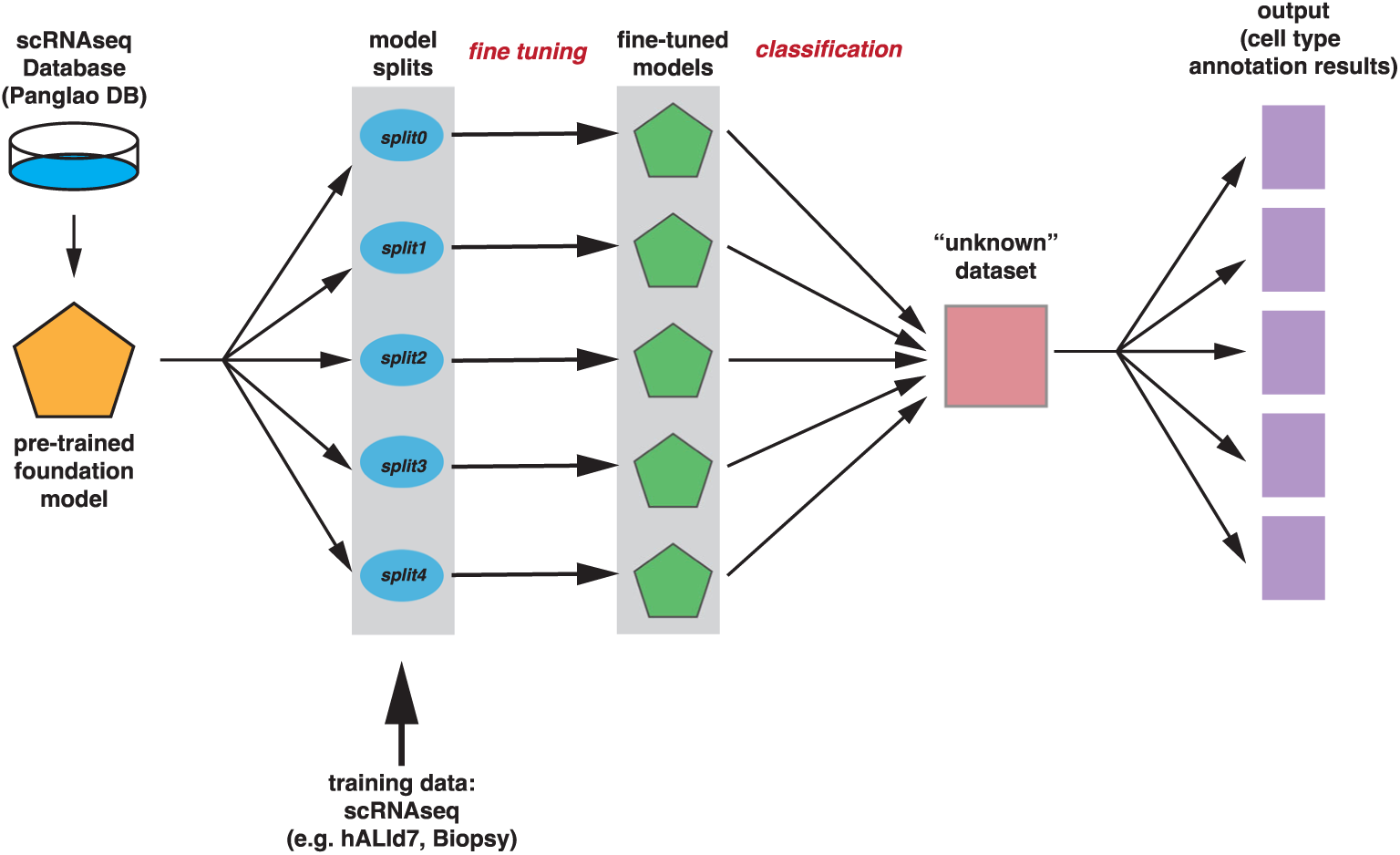
Schematic of BMFM scRNAseq analyses.

**Fig. S6.**
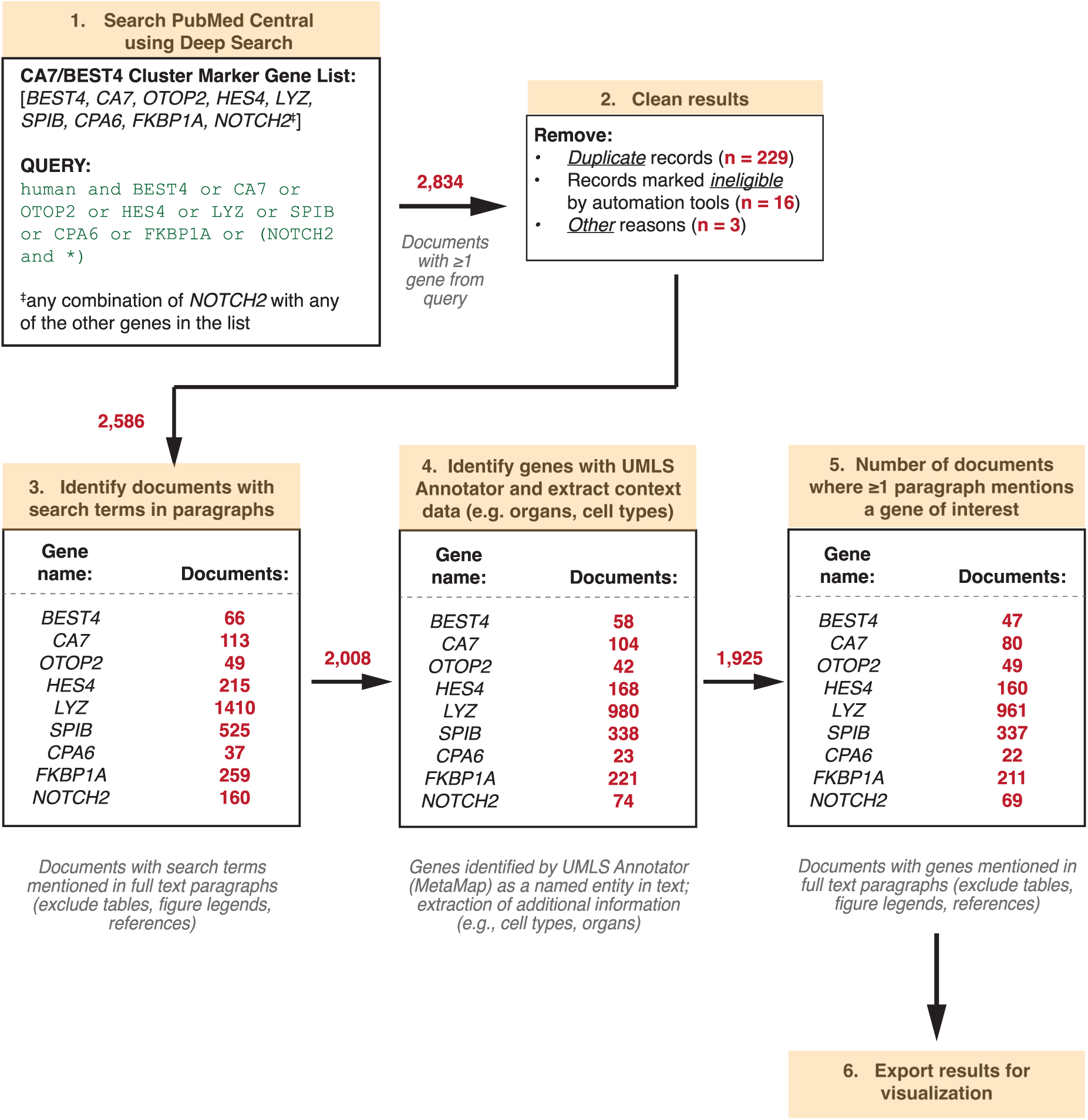
Deep Search Workflow. Workflow for Literature Search and Analysis of BEST4/CA7 Marker Genes using Deep Search (PubMed Central collection).

